# Presence of a SARS-COV-2 protein enhances Amyloid Formation of Serum Amyloid A

**DOI:** 10.1101/2021.05.18.444723

**Authors:** Asis K. Jana, Augustus B. Greenwood, Ulrich H. E. Hansmann

**Affiliations:** Department of Chemistry & Biochemistry, University of Oklahoma, Norman, OK 73019, USA

**Author notes:** Corresponding Author Ulrich H.E. Hansmann - Department of Chemistry & Biochemistry, University of Oklahoma, Norman, Oklahoma 73019, United States.

**Keywords:** Molecular Dynamics Simulations, SARS-COV-2, Serum Amyloid A

## Abstract

A marker for the severeness and disease progress of COVID-19 is overexpression of serum amyloid A (SAA) to levels that in other diseases are associated with a risk for SAA amyloidosis. In order to understand whether SAA amyloidosis could also be a long-term risk of SARS-COV-2 infections we have used long all-atom molecular dynamic simulations to study the effect of a SARS-COV-2 protein segment on SAA amyloid formation. Sampling over 40 µs we find that presence of the nine-residue segment SK9, located at the C-terminus of the Envelope protein, increases the propensity for SAA fibril formation by three mechanisms: it reduces the stability of the lipid-transporting hexamer shifting the equilibrium toward monomers, it increases the frequency of aggregation-prone configurations in the resulting chains, and it raises the stability of SAA fibrils. Our results therefore suggest that SAA amyloidosis and related pathologies may be a long-term risk of SARS-COV-2 infections.

## Introduction

While even after serious complications most COVID-19 survivors appear to recover completely, only little is known about the long-term effects of infections by the severe acute respiratory syndrome coronavirus 2 (SARS-CoV-2). Disease-associated symptoms such as inflammation of blood vessels and overreaction of the immune system^1–3^ are connected with spikes in the concentration of the human Serum Amyloid A (SAA) protein, with the level increasing as the disease progresses from mild to critical^4–6^. The more than thousand times higher blood concentrations of SAA in acute COVID-19 patients^4, 5^ are comparable to the ones seen in patients with various cancers or inflammatory diseases^7^ where the overexpression of SAA is associated with systemic amyloidosis as a secondary illness. SAA amyloidosis is characterized by formation and deposition of SAA amyloids in the blood vessels, causing inflammation, thrombosis and eventually organ damage. Common complication of SAA amyloidosis such as kidney failure or high incidents of thrombosis are also frequently observed in COVID-19 patients^8, 9^. The similarity of symptoms suggests that SAA amyloidosis may exacerbate COVID-19 symptoms^10^, or that it is a long-term risk in COVID-19 survivors causing, for instance, the broad spectrum of symptoms in the multisystem inflammatory syndrome first reported in children and adolescents (MIS-C)^11^, but also observed in adults (MIS-A). This hypothesis is the motivation for the present study where we use molecular dynamics simulations to probe how presence of a SARS-COV-2 protein fragment modulates formation and stability of SAA amyloids. Such SARS-COV-2 triggered amyloid-formation has been observed *in vitro* for *α*Synuclein^12^, but not yet demonstrated for SAA.

The overexpression of SAA in some cancers or inflammatory diseases leads not in all cases to amyloidosis. Usually, after a spike, the concentration levels decrease rapidly in a process that involves dissociation of SAA hexamers followed by cleavage of the released chains. In our previous work^13^, we proposed that the cleavage happens in part because the fragments have a lower probability to re-assemble into the functional hexamers than the complete SAA_1−104_ proteins. We also observed that unlike other fragments, the most commonly found SAA_1−76_ can switch between two structural motifs. The first one is easy to proteolyze (allowing to lower rapidly the SAA concentration) but vulnerable for aggregation, while the opposite is the case for the second motif. If amyloid formation takes longer than proteolysis, the aggregation-prone species dominates. However, if environmental conditions such as low pH encourage amyloid formation, the configurational ensemble shifts toward the more protected form. In this picture, amyloidosis happens when this mechanism for downregulating SAA concentration becomes overwhelmed or otherwise fails. In COVID-19 patients this could happen in three ways: first, presence of SARS-COV-2 could lower the stability of the functional hexamers; secondly, it could increase the probability for association of the SAA fragments after cleavage; and thirdly, it could enhance the stability of the resulting SAA fibrils; each possibility shifting the equilibrium toward SAA fibril formation.

In this computational study, we probe all three possibilities, but focus on the effect of viral proteins, i.e., neglecting the potential roles played by viral RNA. In order to reduce computational costs, we restrict ourselves to short amyloidogenic regions on viral proteins that are most likely to interact with SAA. An example is the nine-residue-segment S^55^FYVYSRVK^63^ (SK9) on the C-terminal tail of the SARS-COV2-Envelope protein, see **Figure 1**. While most of the 75-residue Envelope protein are transmembrane or intracellular, this segment located on the extracellular C-terminal tail. As its location makes it likely to interact with the extracellular SAA proteins, and as its homolog in SARS-COV-1 is known to form amyloids in solution^15^, we investigate in our simulations the interaction of the SK9 segment with the SAA hexamers, monomeric SAA_1-76_- fragments and the SAA fibrils as shown in **Figure 2**. While use of such a small segment may lead to a different mechanism than one would see for the full protein, the danger seems minimal in our cases, as most of the Envelope protein can likely not interact with the extracellular SAA.

**Figure 1:**
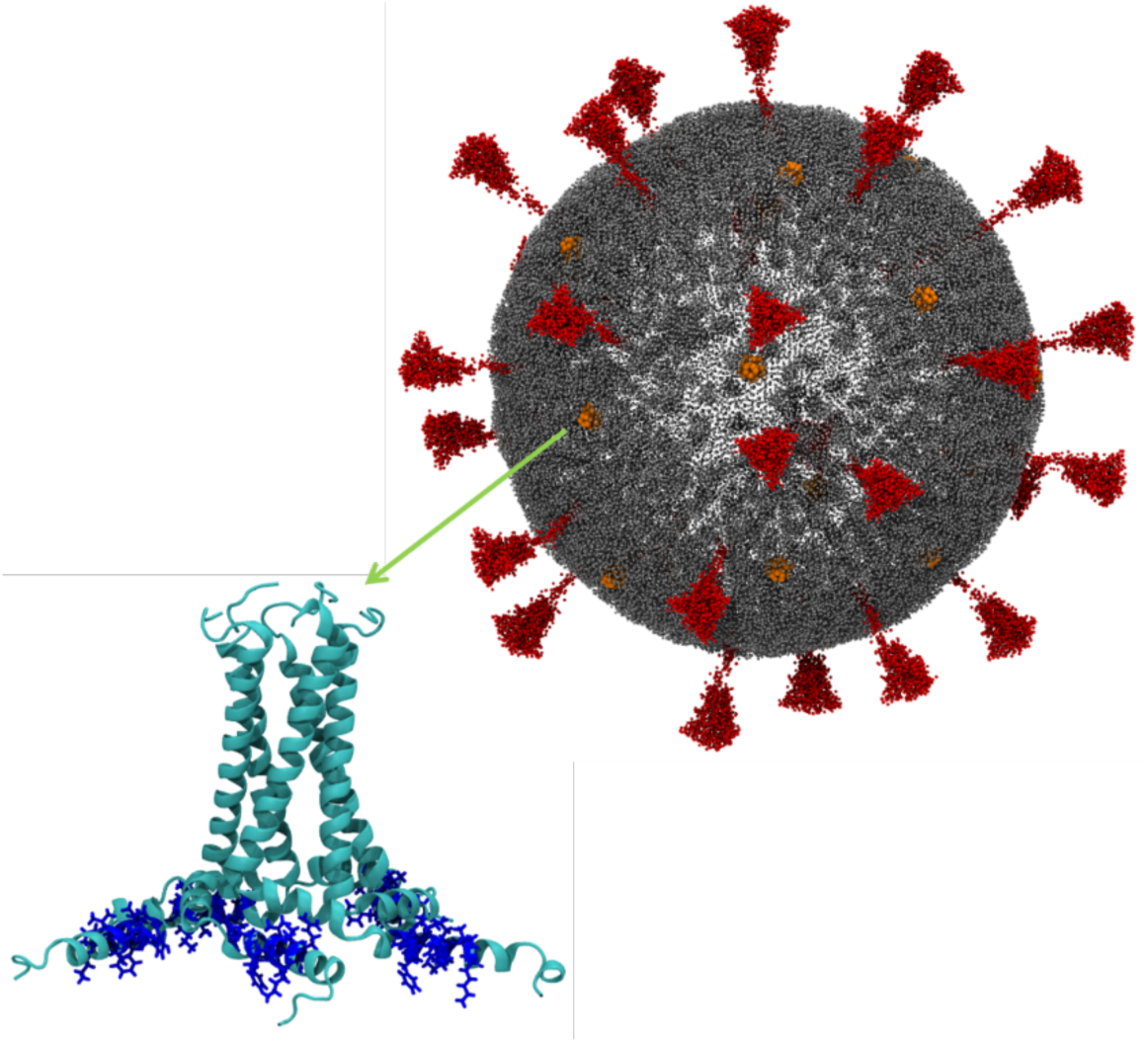
Sketch of the SARS-COV-2 virus with locations of the Envelope (E)-protein marked in orange. Enlarge is also shown the E-protein assembled as a pentamer with the SK9 segments colored in dark blue. The SARS-COV-2 virion is drawn from data in Yu et. al.^14^, while the E-protein pentamer structure is extracted from Ref. 20.

**Figure 2:**
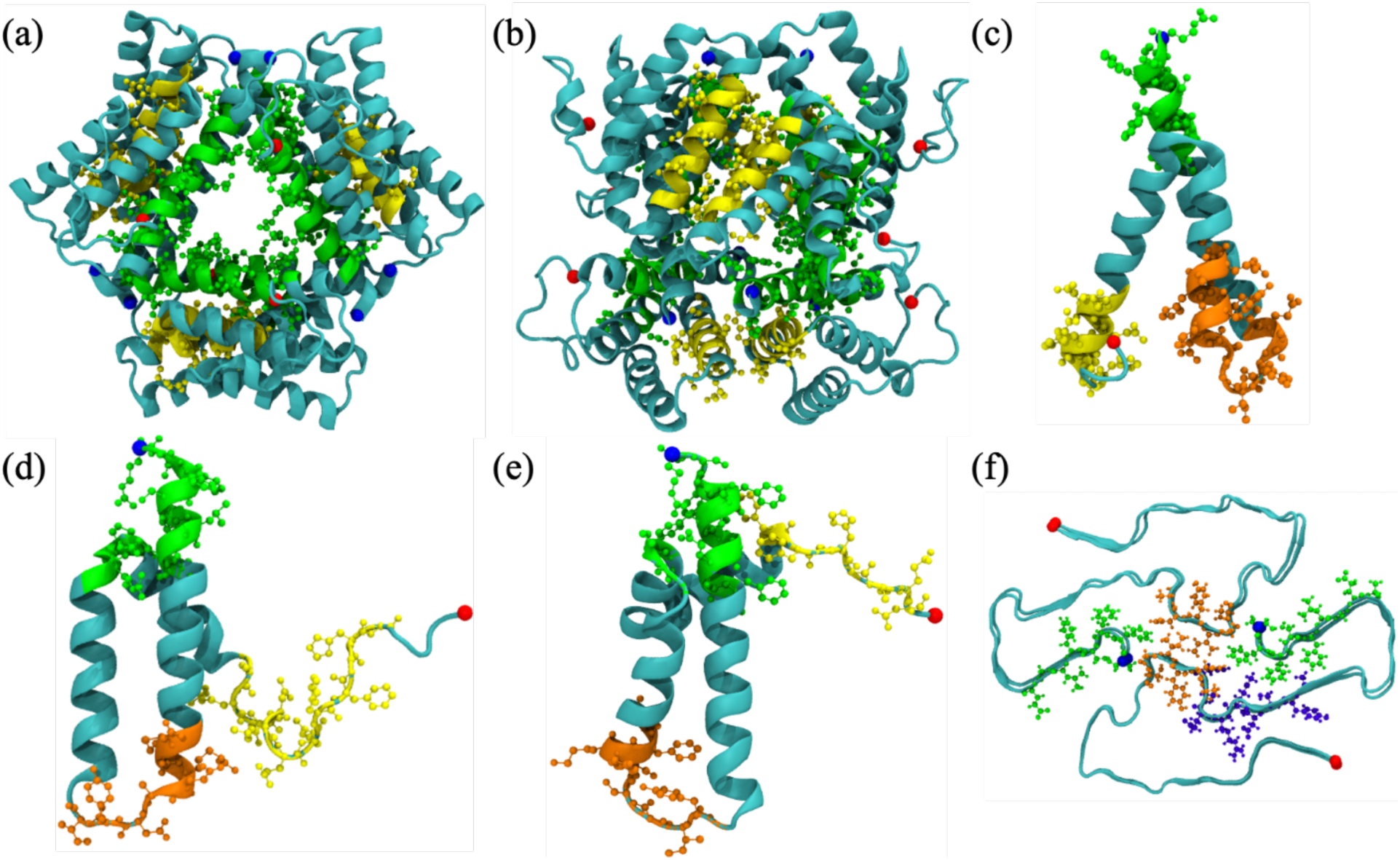
X-ray crystal structure of full-length SAA_1-104_ hexamer (PDB ID: 4IP8) shown in (a) in a top-down view and in (b) in a side view. Each chain consists of four helix-bundles: the N-terminal helix-I (residues 1–27), helix-II (residues 32–47), helix-III (residues 50–69) and the C-terminal helix-IV (residues 73–88). N- and C-terminal residues are here and in all other sub-figures represented by blue and red spheres, respectively. In the start configuration, the virus protein segment SK9 binds with SAA_1-104_ hexamer at the N-terminal helix-I (green) or residues in the helix-III (yellow). The monomer structure of the SAA_1-76_ fragment in (c) is derived from the X-ray crystal structure (PDB-ID: 4IP9), while the representative configurations for the helix-weakened (d) and helix-broken (e) SAA_1-76_ monomers are taken from our previous work^13^. The initial binding positions of SK9 at N-terminal helix-I (green), helix-I-helix-II linker (orange) and disordered C-terminus (yellow) are also shown. Finally, we show in (f) the fibril fragment 2F2L as extracted from the cryo-EM structure of the human SAA fibril (PDB ID: 6MST). In the start configuration, the SK9 segment binds to the SAA_2-55_ fibril at either the N-terminus (green), C-terminal cavity (violet) or the packing interface (orange).

Our results show that presence of viral SK9 fragment raises the risk for SAA fibril formation at all three stages: it reduces the stability of the lipid-transporting hexamer, by increasing the chance for association of the SAA fragments after cleavage, and by enhancing the stability of the resulting SAA fibrils; in each case shifting the equilibrium toward SAA fibril formation. Our results therefore suggest that SAA amyloidosis and related pathologies may be a long-term risk of SARS-COV-2 infections.

## Materials and Methods

### System Preparation

In order to evaluate the effect of the nine-residue SK9- segment S^55^FYVYSRVK^63^ (located in the C-terminal tail of the Envelope protein of the SARS-COV-2 virus) on Serum Amyloid A (SAA) amyloid formation, we evaluate the change in stability of the known hexamer, monomer, and fibril models upon binding, comparing the complex of SK9 and SAA to the corresponding “pure” SAA models.

We choose as the initial configuration for the SAA_1-104_ hexamer the X-ray-resolved crystal structure, deposited in the Protein Data Bank (PDB) under identifier 4IP8^16^ and shown by us in **Figure 2a-b**. Note, that we add here and in the following cases a NH3^+^- group at the N-terminus and a COO^-^ -end group at the C-terminus. These end groups are chosen to make our simulations consistent with our previous work^13^. As *in vivo* SAA monomers are cleaved enzymatically, with SAA_1-76_ the most common resulting fragment, we consider the following three models for our simulation of monomers. The first one, shown in **Figure 2c**, is generated by removing the residues 77−104 from the crystal structure of the full-length SAA monomer (PDB ID: 4IP9)^16^, while the other two configurations were derived by us in Reference 13 as typical motifs found in long-time simulations of the fragment, and named by us “helix-weakened” **(Figure 2d)** and “helix-broken” **(Figure 2e)**. Finally, for simulation of SAA fibrils we have used tetramers made of two folds (protofibrils) and two layers, since we have identified in previous work^17^ such 2F2L tetramers as the smallest stable fibril fragments. Our 2F2L model is shown in **Figure 2f** and is derived from the cryo-EM structure deposited in the PDB database under identifier 6MST^18^. Note that the fibril model is made of SAA fragments with residues 2-55 since no fibril model for human SAA_1-76_ is available. Initially, we built also a second fibril model where we added the likely disordered missing residues 56 to 76 in a configuration predicted by homology modeling. However, we discarded this model for reasons discussed in the result section.

Simulation starting from the above described SAA models serve as a control against which we compare our simulations of the various SAA models interacting with SK9 segments S^55^FYVYSRVK^63^ which is located in the C-terminal tail of the Envelope protein of the SARS-COV-2 virus. As with the exception of the transmembrane domain of residues 8-38^19^ the Envelope protein has not been resolved, we have generated the initial configuration for the SK9 fragment from a model derived by a machine-learning approach and subsequent refinement by molecular dynamics^20^. For this purpose, we have removed from this model the residues 1-54 and 64-75, afterwards capping the remaining nine-residue segment with a NH3^+^ -group at the N-terminus and CONH_2_ as the C-terminus. These end groups were chosen to avoid strong electrostatic interactions between the oppositely charged terminal residues.

Using the AutoDock Vina software^21^, we have generated the start configurations for our simulations by docking the SK9- segments in a ratio 1:1 with the SAA chains in our models. The resulting binding positions of the SK9 segments in the start configurations of either SAA hexamer, monomers, or fibrils, are also shown in **Figure 2**. In the case of the hexamer are the binding positions obtained from a global search, and the configuration with lowest energy is chosen as start configuration. On the other hand, we have performed for each SAA_1−76_ monomer three independent searches, focusing on either the N-terminus, the helix-I – helix-II linker, or the C-terminal region; and chose the respective lowest energy binding pose as the start configuration of the respective runs. Finally, for our fibril models we have again identified SK9 binding positions by a global search using AutoDock Vina.

The various models, either with SK9 binding to the SAA chains or without SK9 present, are each put in a rectangular box with periodic boundary condition. The box is chosen such that there is a minimum distance of 15 Å between any SAA or SK9 atom and all box sides. Each box is filled with TIP3P^22^ water molecules, and counter ions are added to neutralize the system. **Table 1** lists for all systems the number of water molecules and total number of atoms.

**Table 1.**
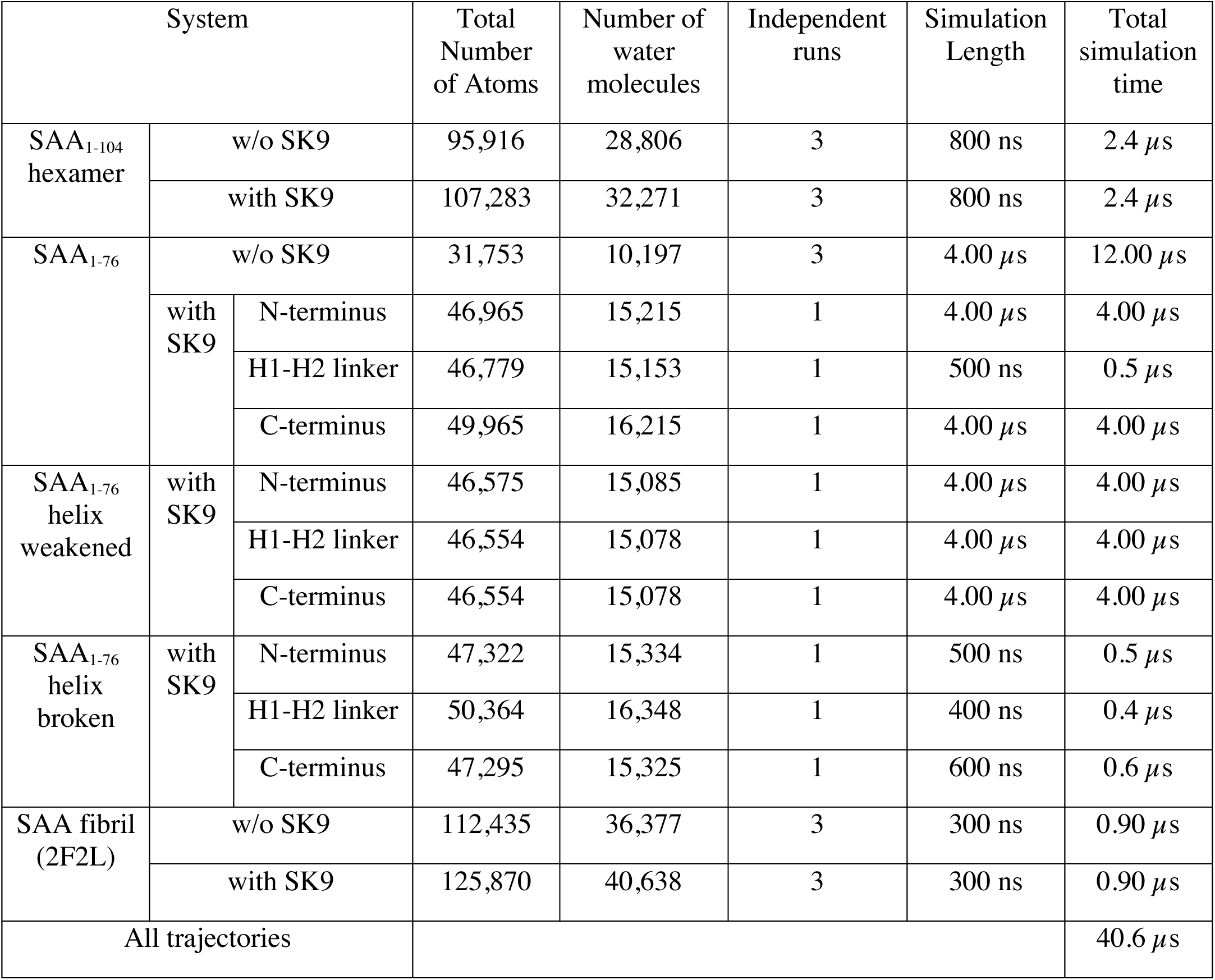
List of Simulated systems and lengths.

| System |  |  | Total<br>Number<br>of Atoms | Number of<br>water<br>molecules | Independent<br>runs | Simulation<br>Length | Total<br>simulation<br>time |
| --- | --- | --- | --- | --- | --- | --- | --- |
| SAA<sub>1-104</sub><br>hexamer | w/o SK9 | | 95,916 | 28,806 | 3 | 800 ns | 2.4 μs |
| | with SK9 | | 107,283 | 32,271 | 3 | 800 ns | 2.4 μs |
| SAA<sub>1-76</sub> | w/o SK9 | | 31,753 | 10,197 | 3 | 4.00 μs | 12.00 μs |
| | with<br>SK9 | N-terminus | 46,965 | 15,215 | 1 | 4.00 μs | 4.00 μs |
| | | H1-H2 linker | 46,779 | 15,153 | 1 | 500 ns | 0.5 μs |
| | | C-terminus | 49,965 | 16,215 | 1 | 4.00 μs | 4.00 μs |
| SAA<sub>1-76</sub><br>helix<br>weakened | with<br>SK9 | N-terminus | 46,575 | 15,085 | 1 | 4.00 μs | 4.00 μs |
| | | H1-H2 linker | 46,554 | 15,078 | 1 | 4.00 μs | 4.00 μs |
| | | C-terminus | 46,554 | 15,078 | 1 | 4.00 μs | 4.00 μs |
| SAA<sub>1-76</sub><br>helix<br>broken | with<br>SK9 | N-terminus | 47,322 | 15,334 | 1 | 500 ns | 0.5 μs |
| | | H1-H2 linker | 50,364 | 16,348 | 1 | 400 ns | 0.4 μs |
| | | C-terminus | 47,295 | 15,325 | 1 | 600 ns | 0.6 μs |
| SAA fibril<br>(2F2L) | w/o SK9 | | 112,435 | 36,377 | 3 | 300 ns | 0.90 μs |
| | with SK9 | | 125,870 | 40,638 | 3 | 300 ns | 0.90 μs |
| All trajectories | | | | | | | 40.6 μs |

### Simulation Protocol

All simulations are carried out using the GROMACS 2018 package^23^ and are employing the CHARMM 36m all-atom force field^24^ and TIP3P water^22^. Number and length of runs are also listed in **Table 1**. For each system, we have taken the configurations generated above, and minimized their energy using the steepest-descent algorithm. This step is followed by 200 ps of molecular dynamic simulations in the NVT ensemble (keeping the volume constant) at 310 K, and a subsequent run of 200 ps in the NPT ensemble keeping the pressure at 1 atm. During the NVT and NPT equilibrations, the nonhydrogen (heavy) atoms of protein are restrained with a force constant of 1,000 kJ mol^-1^ nm^-2^.

The so-equilibrated configurations are the start point of our production runs where the respective systems evolve at a constant temperature of 310 K and a constant pressure of 1 atm. For each set-up we followed three independent trajectories starting from different initial velocity distributions. Note that in these simulations the position of the SK9 segment is not restrained to the original docking sites. Instead, it can move and even detach over the course of the simulation, no longer interacting with SAA. If this happens, the behavior of the system will be the same as for the control (where SK9 is absent). Hence, to save computational resources we stopped a simulation if the SK9 segment did not re-attach within 100ns. The length of all trajectories is listed in **Table 1**.

The temperature is controlled during the simulation with a v-rescale thermostat,^25^ and the pressure with the Parrinello-Rahman barostat,^26^ using a coupling constant of 2 ps. Keeping the water geometry fixed with SETTLE algorithm^27^ and constraining non-water bonds including hydrogen atoms with the LINCS algorithm^28^ allowed us to use a timestep of 2fs for integrating the equations of motion. As we use three-dimensional orthorhombic periodic boundary conditions, we have to use the particle-mesh Ewald (PME)^29^ method for calculating electrostatic interactions. This is done with a real-space cutoff of 12 Å, a value also used as cut-off for Van der Waal interactions, where smoothing started at 10.5 Å.

### Trajectory Analysis

Most of our analysis relies on GROMACS tools such as gmx_rms for calculating the Root-mean-square-deviation (RMSD) with respect to the initial configuration. Another example is the do_dssp tool which implements the Dictionary of Secondary Structure in Proteins (DSSP)^30^ and allows calculation of residue-wise secondary structure propensities. For visualization and for calculating the solvent accessible surface area (SASA) we use VMD software.^31^ Quantities such as the cavity diameter ⟨d_cavity_⟩ in the hexamer are calculated by In-house programs, averaging over the center-of-mass distances between the N-terminal helix-I regions of adjacent units of both layers. Another example is the fraction of native contacts which is calculated using a soft cutoff algorithm and which is defined as^32^:

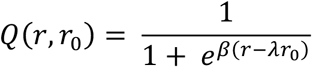

Here, r is the distance between two heavy atoms at which a contact is formed in the native state and r0 denotes this distance in the native state. β denotes smoothing parameter taken to be 5 Å, while λ represents the fluctuation when contact is formed, taken to be 1.8.

Residue-wise binding affinities of the SK9 segment to SAA chains are estimated by calculating binding probabilities instead of free energies. This is because exact methods such as thermodynamic integration would have been too costly for calculating the later, and approximate approaches such as MM/PBSA or MM/GBSA perform poorly when, as in our case, electrostatic interactions between charged (R61 and K63 in the SK9- segment) and polar residues dominate^33^. Here, we define a binding site as the closest residue that has at least one non-hydrogen atom within 4.5 Å from the SK9- segment.

## Results

### Effect of SK9- segments on SAA hexamers

One of the symptoms of COVID-19 is an increase in the concentration of SAA to levels that in a number of cancers and inflammatory diseases often, but not always, leads to amyloidosis as a secondary illness. The response to the overexpression of SAA includes in these diseases a dissociation of the hexamers. The subsequent cleavage of the resulting SAA monomers into fragments is likely because these fragments have a lower probability than the full SAA_1-104_ protein to reassemble as hexamer.^13^ Hence, we start our investigation into the effect of the SK9 segment on SAA amyloidosis by probing how presence of SK9 alters the equilibrium between hexamers and monomers.

As the monomer - hexamer equilibrium depends on the stability of the SAA hexamer, we explore first the change in stability of the SAA hexamer upon binding of SK9 segments. The loss of stability can be monitored by measuring over the length of the trajectory the root-mean-square deviation (RMSD) to the start configuration. If this RMSD is calculated over all heavy atoms in the hexamer, we call this a *global RMSD*. On the other hand, if the RMSD is calculated separately for each chain, and averaged over all six chains, we talk of a chain *RMSD*. Hence, the global RMSD measures the structural deviation of the entire hexamer, whereas the chain-RMSD represents the structural distortion of each chain in the hexamer. Note that we evaluate in both cases the RMSD only for residues 1-76 to stay consistent with our previous work^13^. In **Figure 3a-b**, we plot the time evolution of the global RMSD, averaged over three independent trajectories, comparing the case of SAA bound with the SK9 segment to our control, the hexamer in the absence of SK9. The global RMSD for the SAA hexamer rises in the control simulation initially by nearly 3.0 Å, but quickly reaches a plateau within the first 100 ns, and over the next 400 ns increases only gradually. On the other hand, when the SK9 segments are bound to the SAA chains, the global RMSD rises in the first 100 ns also by about 3.0 Å, but instead of approaching a plateau, continues to rise in a step-wise fashion to a RMSD of 5.0 Å and higher. The differences are much smaller for the chain RMSD, indicating that the interaction with the SK9 segments disturbs the association of SAA chains but not the configuration of the individual chains.

**Figure 3:**
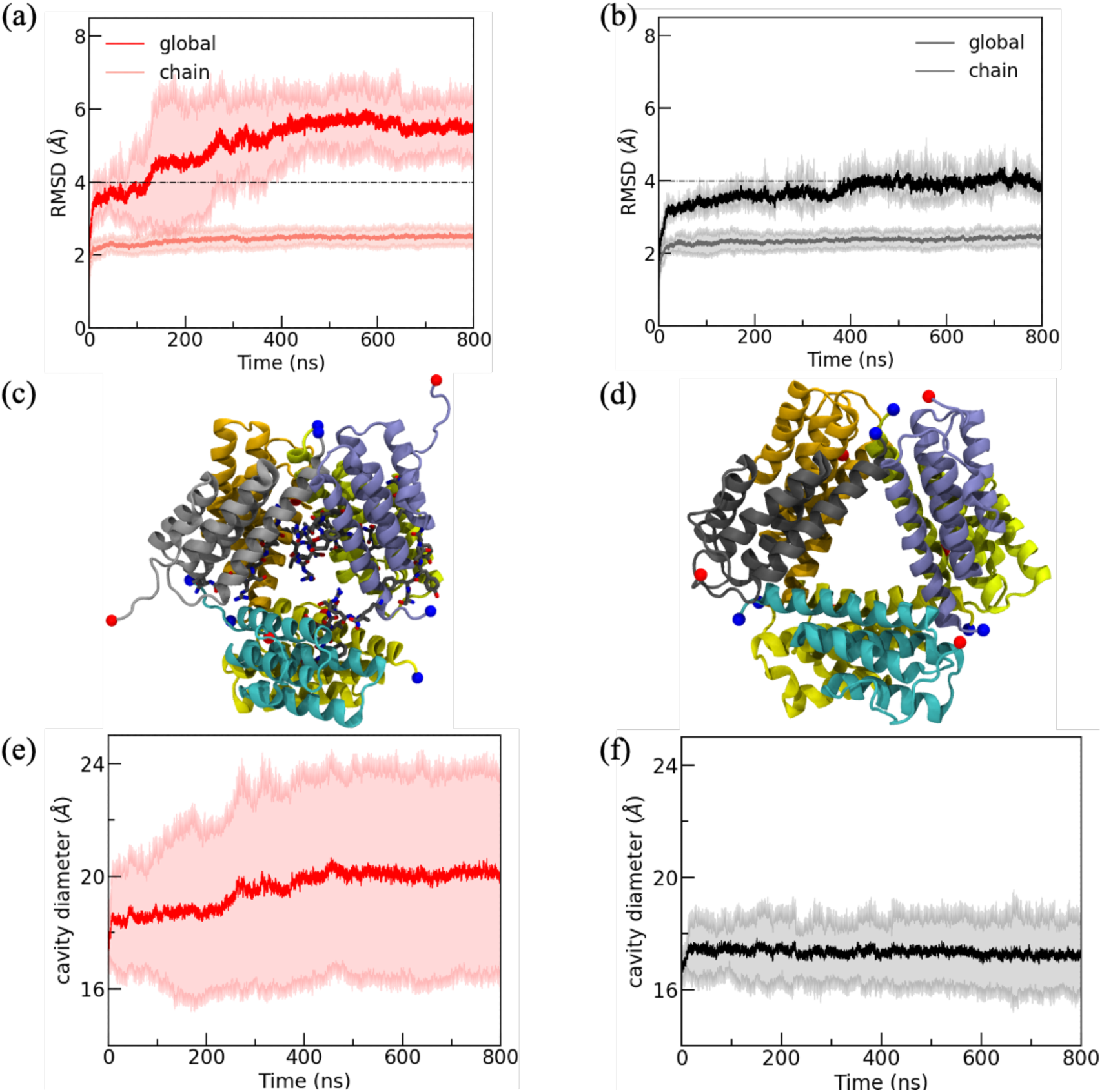
Evolution of the global and chain RMSD of (a) the SAA1-104 hexamer bound with SK9 segments and in (b) our control, the SAA1-104 hexamer in absence of SK9 segments. Representative final configurations (at 800 ns) for the two cases are shown in (c) and (d), respectively, with N- and C-terminal residues represented by blue and red spheres, respectively. Note that the positions of the SK9 segments are also shown in (c). The visual differences are quantified by the cavity diameter plotted in (e) for the SAA1-104 hexamer in complex with SK9, and in (f) for our control. All data are averaged over three trajectories, with the shaded region in (a), (b), and (e), (f) representing the standard deviation over the trajectories.

This can be also seen by comparing the final configurations of both systems in **Figure 3c-d**, where we show the binding of the SK9 segments to the interfacial regions and the resulting distortion of the hexamer. In order to quantify this distortion, we plot in **Figure 3e-f** the time evolution of the cavity diameter ⟨d_cavity_⟩ for both systems. Strongly correlated with the global RMSD we find for the hexamer in absence of SK9 that the cavity diameter approaches quickly a plateau, while for the hexamer in complex with SK9 segments the values again increase stepwise, with a pronounced step at in <d_cavity_> at 250 ns.

In order to understand in more detail how SK9 affects the association of the SAA chains in the hexamer, we first calculate the residue-wise binding probability of SK9- segment towards SAA hexamer. Data are averaged over the final 500 ns of each trajectory and shown in **Figure 4.** The binding probability map demonstrates that SK9- segment preferentially binds with interfacial aromatic and hydrophobic residues (F3, F4, W18 and I65), and inter-chain salt-bridge forming residues (D12 and R25), thereby disrupting the interfacial hydrophobic and stacking contacts, as well as inter-chain salt-bridges. As a consequence, the solvent-exposure of the hydrophobic residues in the hexamer increases. This can be seen in **Figure 5a** where we show the solvent-accessible surface area (SASA) of exposed hydrophobic residues, measured in each of the three trajectories over the last 500 ns using VMD with a spherical probe of 1.4 Å radius. We find that in the presence of SK9 the SASA distribution is shifted towards higher values indicating solvent exposure of interfacial residues. The mean SASA value for hydrophobic residues in presence of SK9 is (10,320 ± 418) Å^2^, and (10,426 ± 260) Å^2^ in absence of SK9. Thus the hydrophobic solvent accessible surface increases by about 9 % in presence of SK9 segments, leading to the lower stability of the hexamer indicated by the higher global RMSD values in **Figure 3a**.

**Figure 4:**
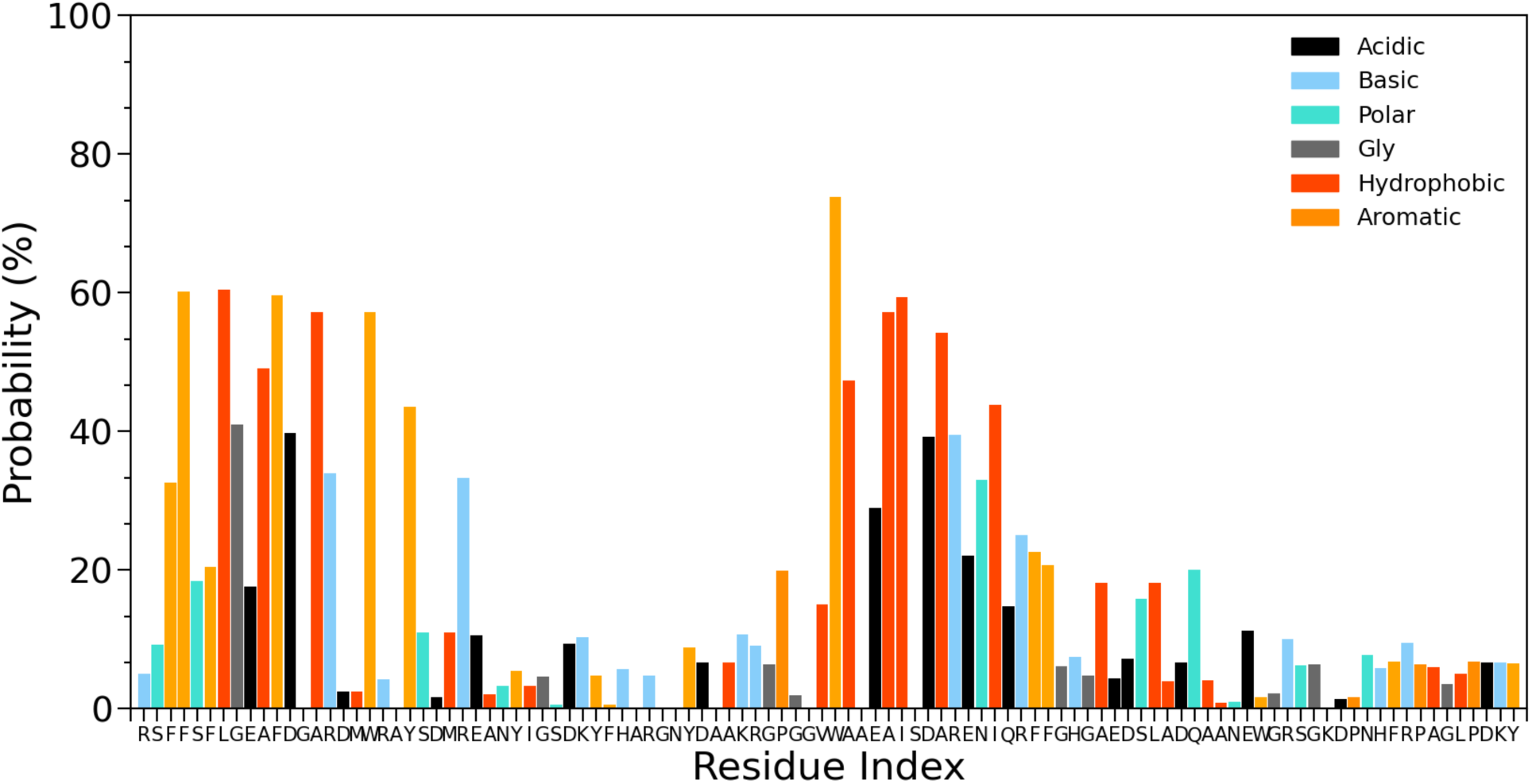
Residue-wise binding probability of SK9- segment towards full-length SAA_1-104_ hexamer. Data are averaged over the final 500 ns of each trajectory.

**Figure 5:**
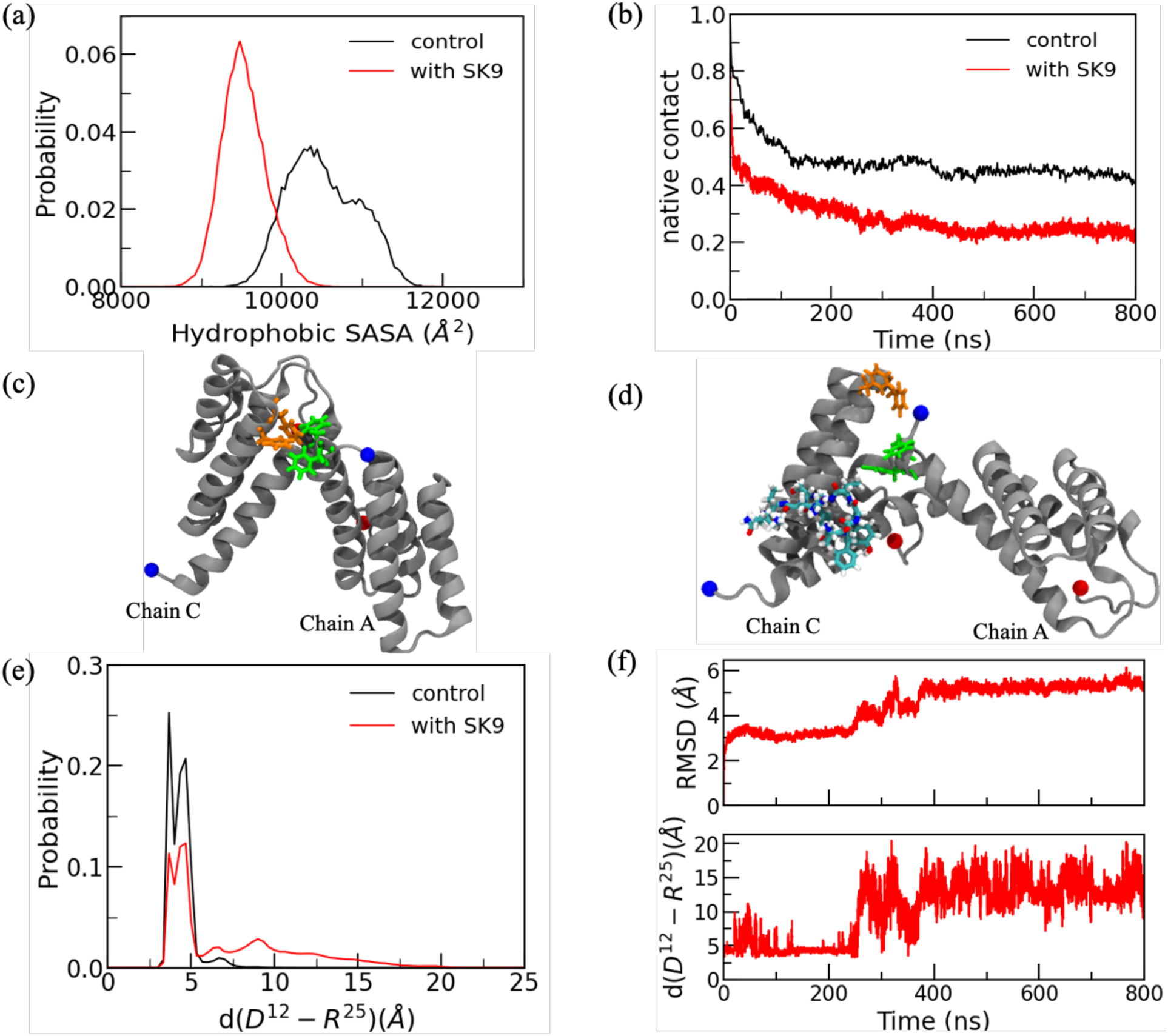
(a) Normalized distributions of the solvent accessible surface area (SASA) of hydrophobic residues. (b) Fraction of the inter-chain native contacts as a function of time, averaged over all six chains in a hexamer and all three trajectories. Representative snapshot of the A and C chain, extracted from a SAA hexamer in absence (c) and presence (d) of SK9, with N- and C- terminal residues represented by blue and red spheres, respectively. The residues F^3^ and F^4^ from chain A, and F^68^ and F^69^ from chain C and ligand are in licorice representation. The SK9 segment is drawn in cyan. (e) Normalized distribution of D^12^-R^25^ salt-bridge distance. (f) Time evolution of the global RMSD and the center of mass distance between D^12^ of chain A and R^25^ of chain C, both measured in Å, along a representative trajectory of SK9- bound SAA_1-104_ hexamer.

The loss in contacts resulting from binding to the SK9 segment can be seen in **Figure 5b** where we plot the time evolution of the number of native inter-chain contacts, i.e., contacts between SAA chains in the hexamer that also exist in the start configuration. We define contacts by a distance cut-off of 4.5 Å and calculate the fraction of the inter-chain native contacts as described in the method section. In both systems decreases this fraction within the first 50 ns. However, it reaches a stable value with little fluctuation after 100 ns in the control simulation, whereas in presence of SK9 the loss of contacts continues and is clearly more pronounced after about 250ns. For a more fine-grained picture we have calculated the inter-residue contact probabilities between monomers; defining two residues whose heavy atoms lie within 7 Å of each other as a contact pair. Analyzing these contacts, we find multiple π-π stacking and hydrophobic interactions involving the residues F^3^, F^4^, W^18^, F^68^, F^69^ and I^65^ that stabilize the interface region, as do the inter-chain salt bridges between pairs D^12^-R^25^, E^9^-R^25^ and E^26^-R^47^. On average we find in our control, that is for the hexamer in absence of the SK9 segments, (1078± 96) interfacial contacts. However, when present, the SK9 segments bind with the above listed interfacial residues, disrupting the interchain network in the hexamer, and the number of interfacial contacts decreases to an average of (704± 81) contacts. As an example, we show in **Figure 5c-d** representative snapshots of the A and C chain from the control simulation and from the hexamer binding with SK9. In the control simulation (**Figure 5c**) are residues F^3^ and F^4^ from chain A in contact with residues F^68^ and F^69^ from chain C, while in **Figure 5d** tyrosine and valine residues from a SK9 segment bind with the residues F^3^ and F^4^ from chain A, thereby preventing π-π stacking with residues F^68^ and F^69^ from chain C. We have quantified this effect for the salt bridge forming residues D^12^ and R^25^. For this pair, we plot in **Figure 5e** the probability distribution of the center of mass distance for both the control, and for the hexamer in presence of SK9 segments. Measurements for both systems are taken over the last 500 ns of all three respective trajectories. The shift in the center-of-mass distance toward larger values in presence of SK9 demonstrates the disruption of the interfacial salt bridge formed by the two residues D^12^ and R^25^ in the control. The loss of this salt bridge in presence of SK9 destabilizes the SAA hexamer. This can be seen in **Figure 5f** where for a representative trajectory of the SAA hexamer in presence of SK9 segments we compare the time evolution of the center of mass distance between D^12^ of chain A and R^25^ of chain C with the time evolution of the global RMSD. The strong correlation between both quantities is especially visible at 250 ns where the loss of the salt bridge (once the distance between the two residues is larger than 4.0 Å) is mirrored by a jump in RMSD.

Hence, our first result is that the binding of SK9 segment with the SAA hexamer competes with the inter-chain binding of the SAA proteins, reducing the stability of the hexamer. Our computational resources did not allow us to observe disassembly of the hexamer, but the loss of stability indicates that in presence of the SK9 segments the equilibrium is shifted away from the hexamer and toward monomers. While this shift downregulates the activity of SAA, it also increases the chance for aggregation.

### Effect of SK9- segments on SAA_1-76_ monomers

We argue in Ref. 13 that after dissociation of the hexamer the monomers are cleaved into fragments because this eases their proteolysis. The most common fragment SAA_1-76_ is special in that it evolves into an ensemble of configurations dominated by two forms. The first one is easy to proteolyze (allowing for a quick decrease in SAA concentration) but vulnerable to aggregation, while the situation is reversed for the second motif. If amyloid formation takes longer than proteolysis, the aggregation-prone species prevails. However, if external factors encourage amyloid formation, the frequency of the more protected motif increases. Hence, in the second part of our investigation we study how the interaction with SK9 alters this mechanism.

For this purpose, we have simulated three systems. The first one is a SAA_1-76_-fragment in the same configuration as seen in the full-sized free monomer (PDB-ID: 4IP9). This is presumably the structure of the fragment right after cleavage. We follow the time evolution of this fragment in the presence of SK9 segments in three molecular dynamics simulations, with the SK9 segment initially docked to either the helix-I – helix-II linker, the N-terminus, or to the C-terminal region. We then compare our results from these three simulations with control simulations where the SK9- segment is absent. When initially binding to the helix-I - helix-II linker, the SK9 segment separates within 500 ns from the SAA monomer and moves away, easily monitored by the time evolution of the number of contacts between SK9 and the SAA monomer, a quantity that at separation drops from fluctuating between 100 and 300 to zero. As after separation the time evolution of the SAA chain is similar to the control simulation, we discuss in the following only the other two cases. For instance, we show in **Figure 6** the time evolution of the secondary structure for the trajectory where the SK9- segment initially binds with the disordered C-terminal region. Over the course of the simulation the SK9 segment shifts its binding partners to hydrophobic residues in N-terminal helix-I (L^7^, A^10^ and F^11^) and some residues encompassing the helix-III and disordered C-terminal region (W^53^, A^54^, E^56^, A^57^, D^60^, N^64^, I^65^, R^67^, F^68^, G^70^ and E^74^), see also the corresponding snapshot in **Figure 6**. As a consequence of this binding pattern, we observe formation of a β-hairpin, involving residues 28–30 and 33–35 in the helix-I – helix-II linker and in the beginning of helix-III. Helix-II (residues 32-47) unfolds completely over the course of the simulation. This β-hairpin formation was also observed in a previous study by Gursky et. Al,^34^ but is on the time scale of our simulations not seen in the control simulations. Both the unfolding of helix-I and helix-II and the β-hairpin formation in the helix-I – helix-II linker region are also not observed, when the SK9- segment binds initially to the N-terminal helix-I, but the two trajectories are otherwise similar. Configurations in both trajectories resemble the easy-to-proteolyze, but aggregation prone, helix-weakened configurations of **Figure 2d,** sharing similar reduced helicity of residues 63-69 in helix-III (the mean helicity percentages are (43 ±41)% and (69 ±31)%, respectively) and a comparable solvent accessible surface area for the first eleven N-terminal residues (with mean SASA values of (524 ±90) Å^2^, and (552 ±43) Å^2^ respectively).

**Figure 6:**
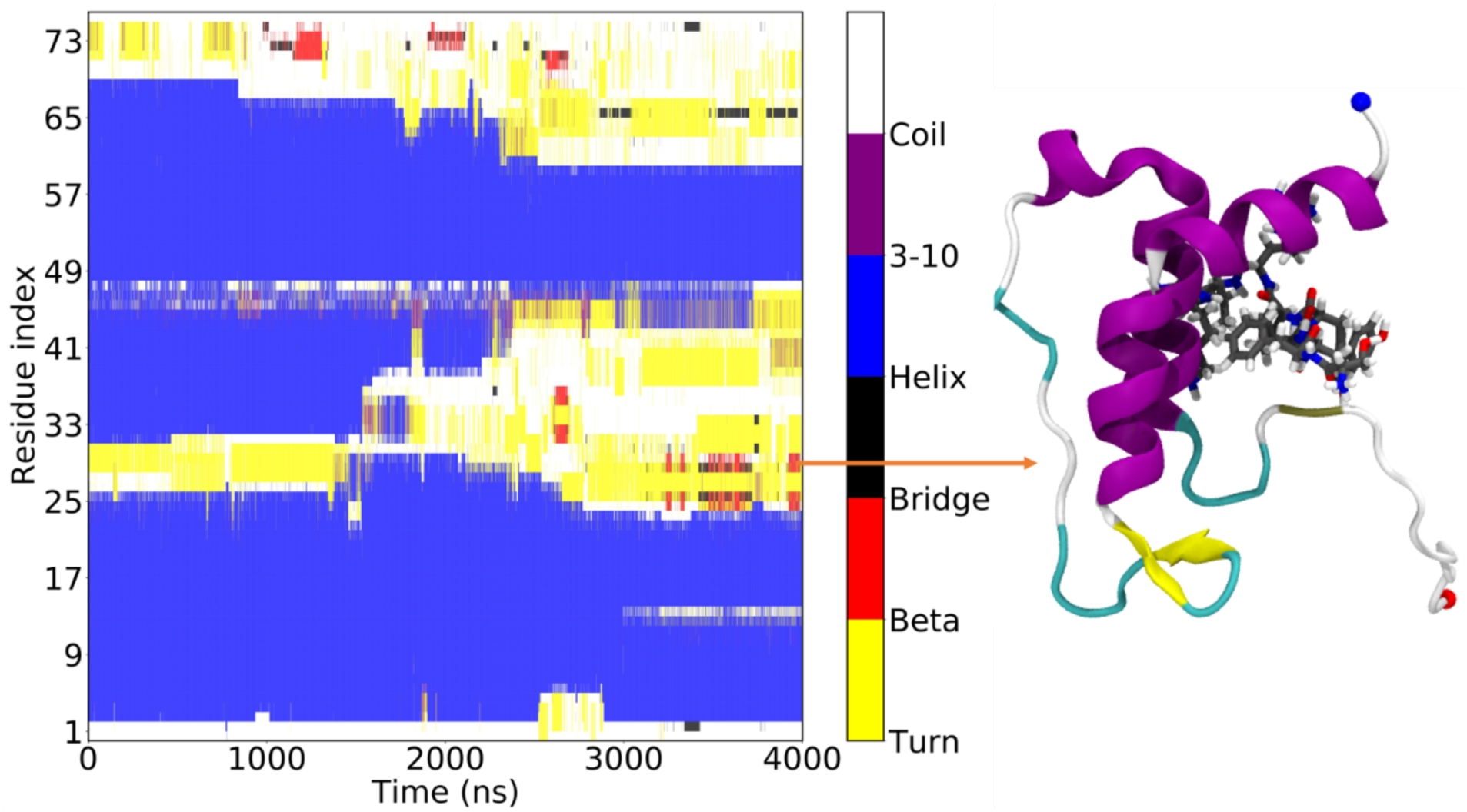
Residue-wise secondary structure map along a SAA_1-76_ monomer simulation in presence of SK9. The initial SAA_1-76_ monomer structure is extracted from the X-ray crystal structure (PDB-ID: 4IP9) with N- and C-terminal residues marked by blue and red spheres, respectively.

Without the viral amyloidogenic segment present, the SAA_1-76_-fragment can also evolve into the helix-broken form of **Figure 2e** which is more difficult to proteolyze but also less aggregation-prone. Our simulations indicate that presence of SK9 alters the distribution of the two forms, shifting it to the more aggregation-prone helix-weakened form. Depending on the effect that presence of SK9 has on the stability of the two forms this would imply a raised probability for SAA-fibril formation. Hence, we have also performed molecular dynamic simulations of helix-broken and helix weakened SAA_1-76_-fragments interacting with a SK9 segment initially docked to either the N-terminus, the helix-I – helix-I -linker, or the C-terminal region.

Independent on where the SK9 segment is initially docked, we observe that in all our trajectories, starting from SAA_1-76_ in the helix-broken configuration, the virus segment disengages and moves away. Consequently, no noticeable differences to the control simulation are seen. This observation suggests that once the SAA fragment assumes a helix-broken configuration, it will not be affected by presence of the viral segment and will stay in this less aggregation-prone motif. On the other hand, when the SK9 segment is initially bound to SAA_1-76_ in the helix-weakened form, we find after about 800 ns a clear signal for forming a N-terminal *β*-strand involving the first eleven residues. This observation is independent on where the SK9 segment initially binds to the SAA chain, see, for instance, the snapshots of representative configurations in **Figure 7a.** The appearance of a N-terminal *β*-strand is important as this region is known to be crucial for amyloid formation^35–37^, and we have shown in Ref. 13 that fibril assembly starts with this region.

**Figure 7:**
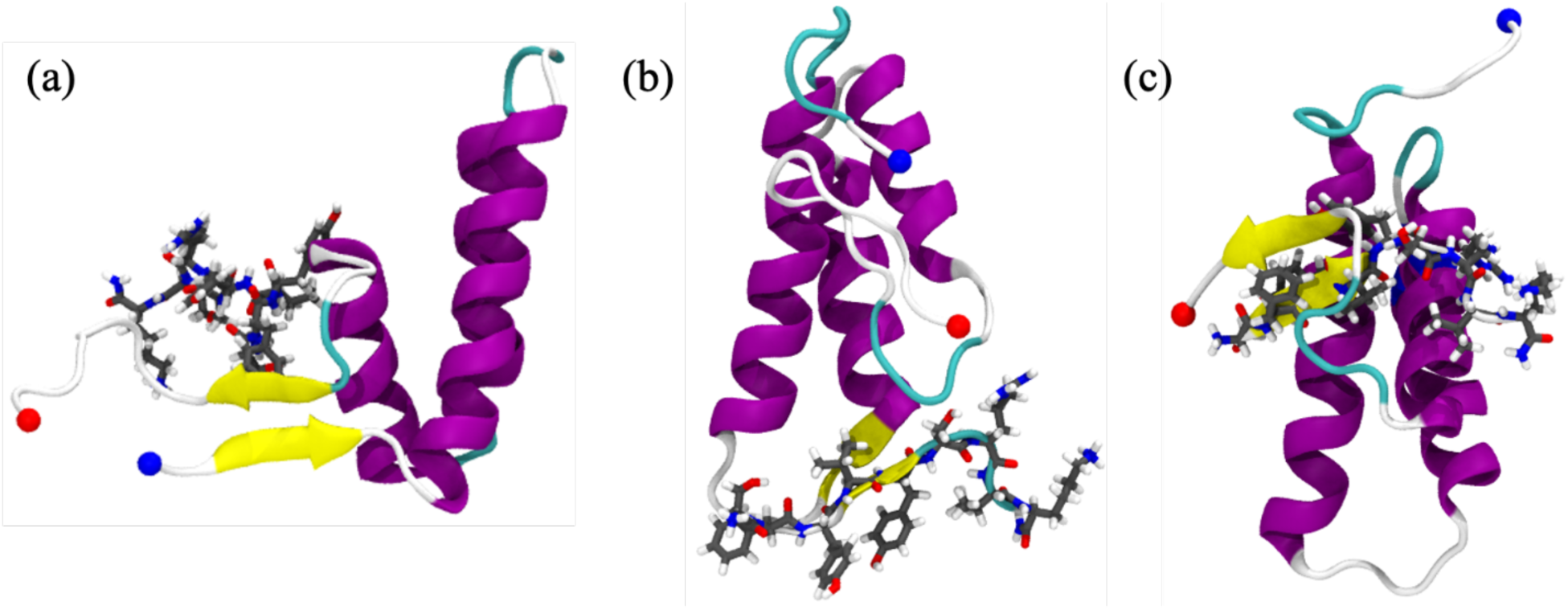
Representative snapshots extracted from a simulation of the SAA1-76 monomer in the presence of SK9, binding to either the N-terminus (a), the helix-II – helix-III linker (b), or the C-terminus (c). In the start configuration the SAA fragment is in all three trajectories in the helix-weakened form. N-and C-terminus are marked by a blue and red sphere, respectively.

As in the earlier discussed simulations starting from a native-like configuration for the SAA_1-76_ fragment, we also observe the β -sheet formation in the helix-I - helix-II linker region or the disordered C-terminal region, see the corresponding snapshots in **Figure 7b** and **7c**. Note that while the initial binding sites are obtained from docking calculations, visual inspection of the trajectories shows that the SK9 segment does not stay bound at the initial position. For this reason, we have also calculated the residue-wise binding probabilities of SK9- segment towards the helix-weakened SAA_1-76_ conformations. Data are averaged over the final 3.0 *µ*s of each trajectory and shown in **Figure 8.** These binding probabilities indicate that the SK9 segment binds preferentially with hydrophobic (A^10^ and A^14^) and aromatic residues (F^11^ and W^18^) in N-terminal helix-I region, along with binding to helix-III and the disordered C-terminal region through hydrophobic (A^54^, A^57^, I^58^ and A^61^), aromatic (W^53^, F^68^ and F^69^) and electrostatic interaction (E^56^, D^60^, E^74^ and D^75^). Our data indicate that the binding affinity of the SK9- segment towards the disordered C-terminus is stronger than to the N-terminus. Note that the binding probability of SK9- segment towards the helix-I - helix-II linker region is relatively low in our simulations, but when binding occurs, the segment forms of a β-hairpin with the linker region (shown in **Figure 7b**).

**Figure 8:**
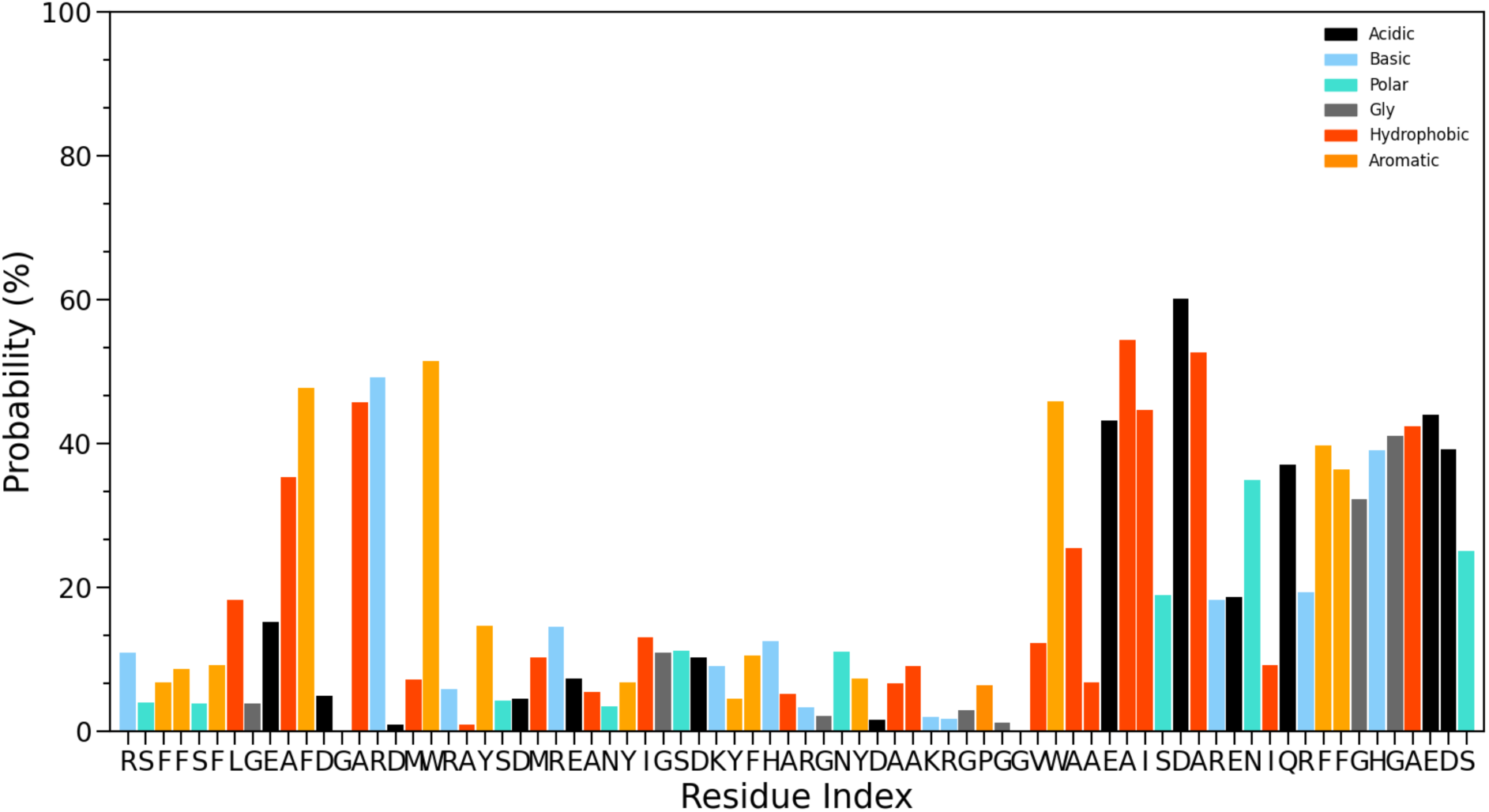
Residue-wise normalized binding probability of SK9- segment for the helix-weakened SAA1-76 conformations. Data were averaged over the final 3.00 µs of each trajectory.

The increased propensity for β-strand formation is confirmed by **Figure 9** where we plot the secondary structure as function of time. No β-strands are formed in the control simulations (**Figure 7a**), while there is a clear signal for it in **Figure 7b** where we show the same quantity for a trajectory where the SAA monomer binds initially with the SK9 segment at the helix-I - helix-II linker region.

**Figure 9:**
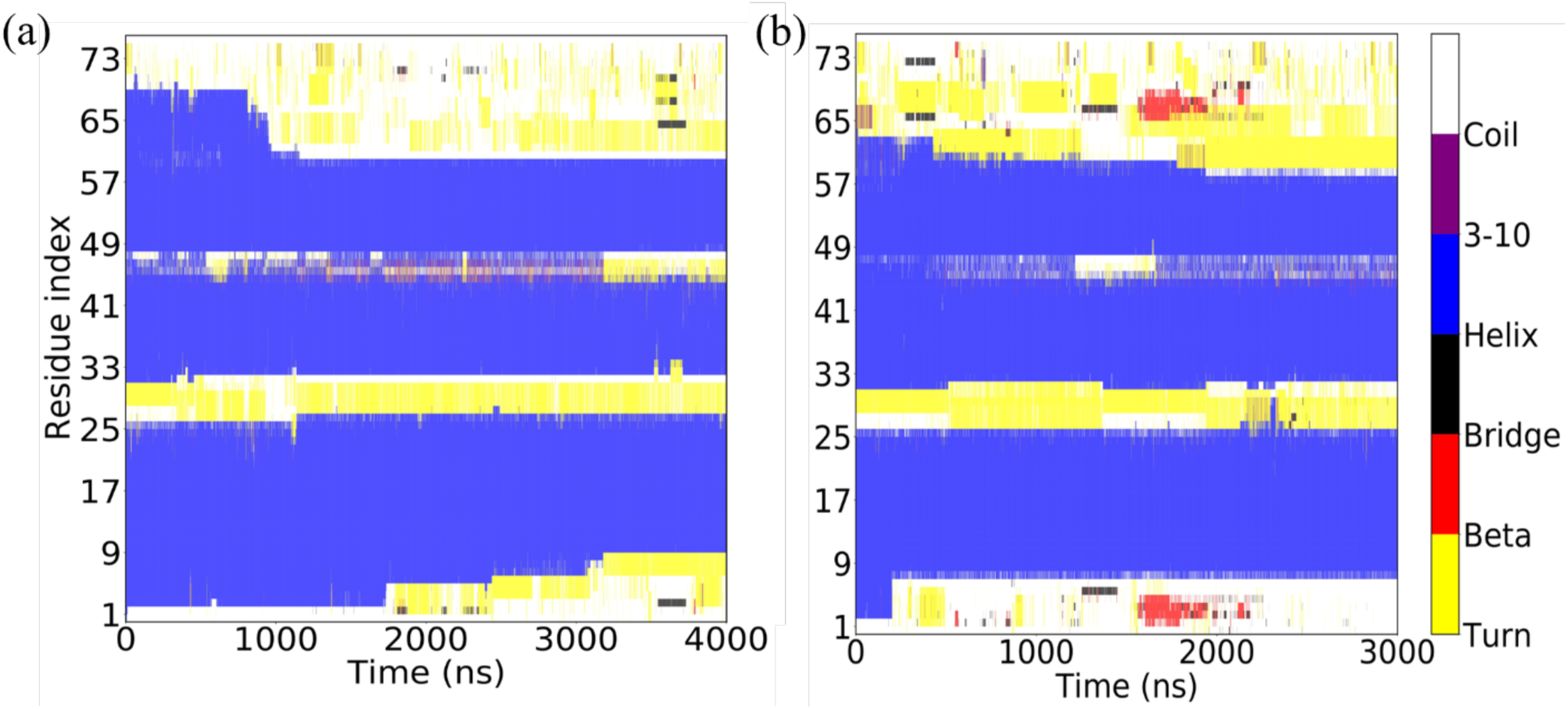
Residue-wise secondary structure measured along a trajectory of the SAA1-76 monomer in (a) absence and (b) presence of the SK9- segment. Both simulations start with the SAA fragment in a helix-weakened configuration.

Hence, a second effect by which SK9 can increase SAA amyloid formation is by shifting the equilibrium toward helix-weakened SAA_1-76_-fragments and by initiating the β-sheet formation in these fragments. The higher probability for forming an N-terminal *β*-strand (residues S^2^-S^5^), known to be the start point for fibril formation^35, 36, 38^, results from interaction between N-terminus (residues 1-6) and C-terminal region (residues 65-72), more precisely π-π stacking (F^3^-F^68^ and F^4^-F^69^) and hydrophobic (F^6^-I^65^) interactions. In order to specify these relations we show in **Figure 10** the difference between the residue-residue contact probabilities which measured in simulations of the helix-weakened SAA_1-76_ in the presence of a SK9- segment, and the ones measured in the control simulations where no SK9 segment is present. The contact propensities are averaged over the last 3.0 µs of all three trajectories. These relative contact probabilities show that hydrophobic interaction between helix-I and helix-III involving residues A^10^-A^54^, A^14^-I^58^, M^17^-I^58^ and Y^21^-R^62^ are significantly enhanced in the presence of the SK9- segment. In contrast, the SK9- segment weakens the interactions within the N-terminal helix-I (F^4^-A^14^ and S^5^-F^11^). SK9 binding also increases the salt-bridge propensity of E^9^-R^47^ and decreases the salt-bridge propensity of D^16^-R^47^. Of special interest is the change in the contact pattern in the helix-I - helix-II linker region. Here, the SK9- segment increases the hydrophobic interactions between the helix-I - helix-II linker region and the end of helix-I involving the residue pairs M^24^-I^30^, M^24^-F^36^ and A^20^-F^36^, but decreases the hydrophobic interactions between helix-II or helix-III and the disordered C-terminal region involving residues pairs F^36^-A^55^ and F^36^-I^58^. Hence, the SK9– segment disrupts residue-residue contact pattern in the otherwise unchanged helix-I and helix-III region, thereby inducing the β-sheet formation in the N-terminal helix-I and the helix-I - helix-II linker region.

**Figure 10:**
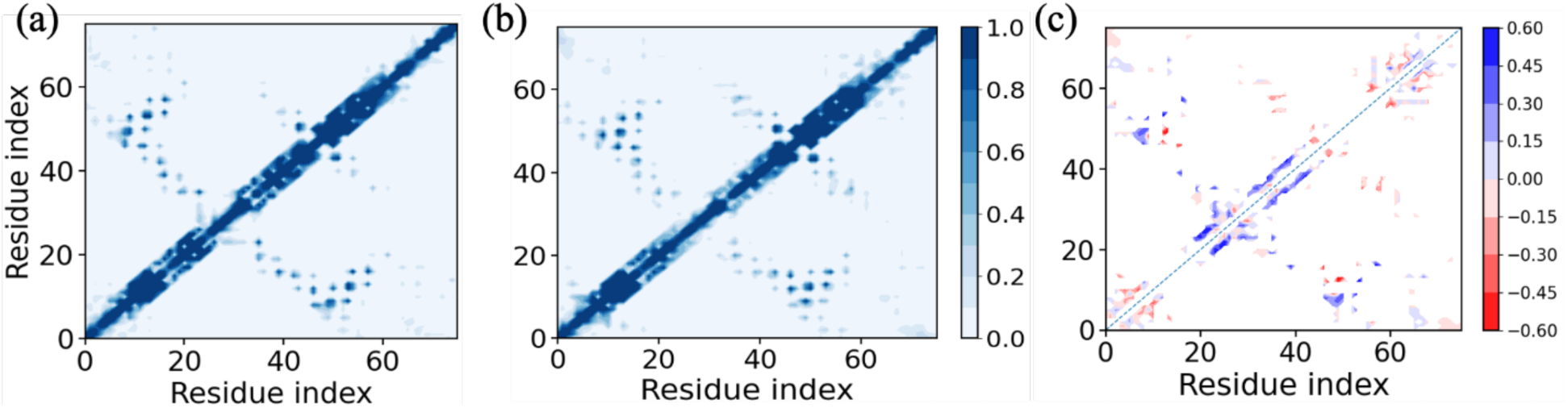
Residue-residue contact probabilities of helix-weakened SAA1-76 conformations in (a) the presence and (b) in the absence of SK9- segment. The difference between the two quantities is shown in (c). Data were averaged over the final 3µs of each trajectory.

### Effect of SK9- segments on the stability of SAA fibrils

In the previous two sections we have shown that the SK9 segments reduce the stability of the hexamer, making it easier to dissolve the hexamer; and that they raise the aggregation propensity of the monomer fragments by encouraging the N-terminal residues to assume a *β*-strand configuration that is known to be crucial for fibril formation^35–38^. In our third set of simulations, we finally study the effect of the amyloidogenic segment on the stability of SAA fibrils. This is because stabilization of the fibril will shift the equilibrium between hexamers, monomers, and fibrils toward the fibrils. Having identified in previous work^17^ a two-fold-two-layer (2F2L) tetramer as the critical size for fibril stability, we have simulated the effect of SK9 on such tetramers. The creation of the start configuration and the set-up of the simulations are described in the method section. Note that our fibril model is made of segments SAA_2-55_ as no fibril model for human SAA_1-76_ is available. A second fibril model with the missing residues 56 to 76 added by homology modeling was no longer considered by us after we realized that the SK9 segment does not bind to the disordered region of residues 56-76. This is because we had seen in previous work^17^ that the added presence of the residues 56-76 by itself does not affect the stability of the SAA_2-55_ chains in the fibril form. Hence, we have only performed molecular dynamics simulations of the 2F2L tetramer, built from SAA_2-55_ chains, either in the presence or -as control – in the absence of SK9 segments.

Following the two systems over 300 ns, we compare the change in stability along the trajectories between the two cases. Representative final configurations are shown together with the corresponding start configurations in **Figure 11**. Visual inspection of these configurations suggests that the fibril is stabilized by the SK9 segment.

**Figure 11:**
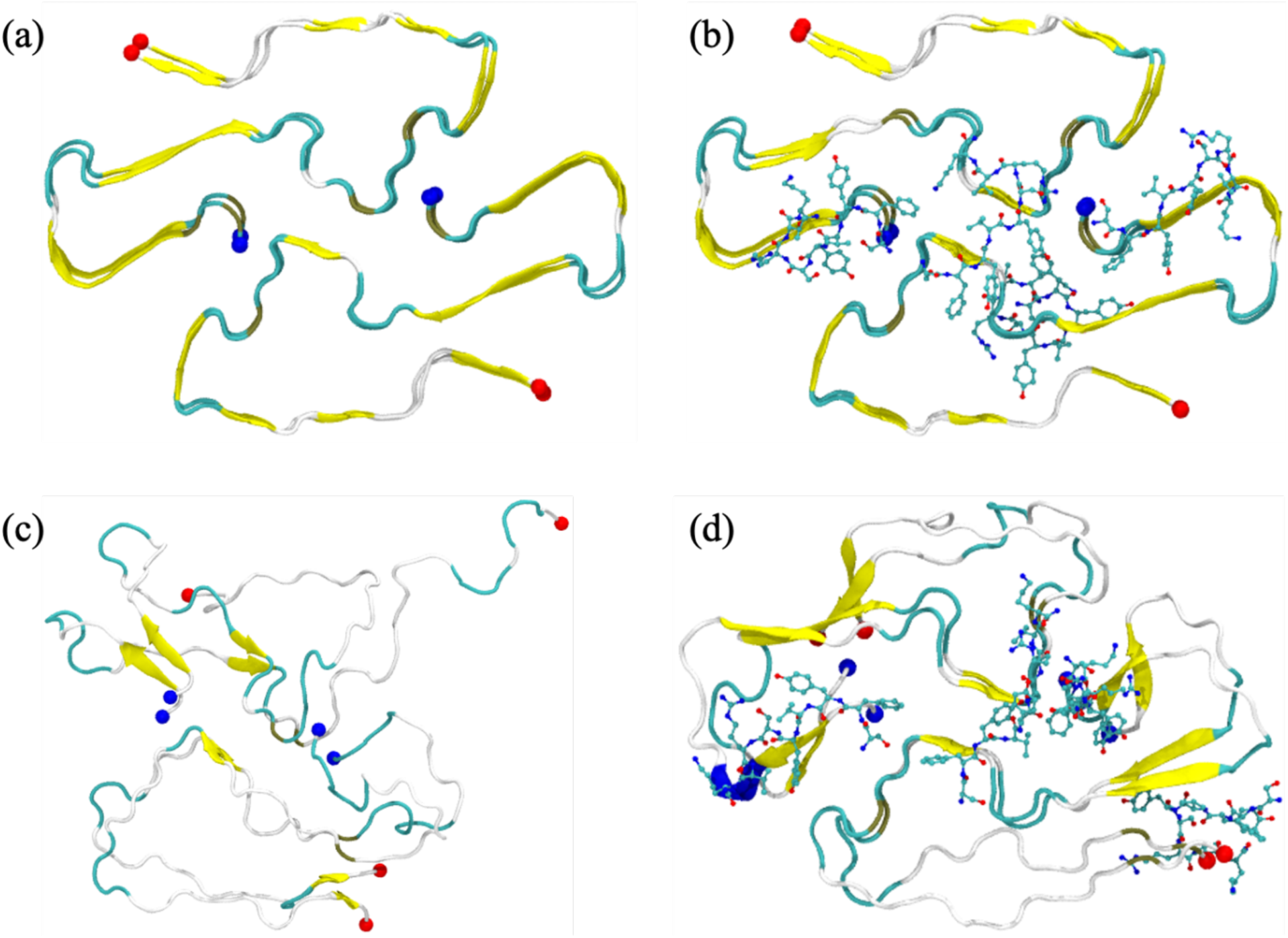
The initial configuration of our control simulation, the SAA_2-55_ fibril in absence of SK9 segments, is shown in (a), while the SAA_2-55_ fibril bound with SK9 segments is shown in (b). Representative final configurations (at 300 ns) for the two cases are shown in (c) and (d). N- and C-terminal residues are represented by blue and red spheres, respectively.

The above visual inspection is again quantified by following the time evolution of global RMSD (measured over the whole fibril) and chain RMSD in **Figure 12**, where the later one is the average over RMSD measurements of the individual chains. Similar to the hexamer, we observe for both systems an initial increase in both RMSD values, which plateaus for the SK9- bound fibril after about 100 ns while the RMSD continues to rise for the fibril in absence of the viral segment until reaching a global RMSD of 8.0 Å after 200 ns. The stabilization of the SAA fibril in presence of SK9 is also supported by the smaller root-mean-square-fluctuation, also shown in **Figure 12** and calculated over the last 200 ns, as they indicate that the shape of the individual chains changes less than in the control.

**Figure 12:**
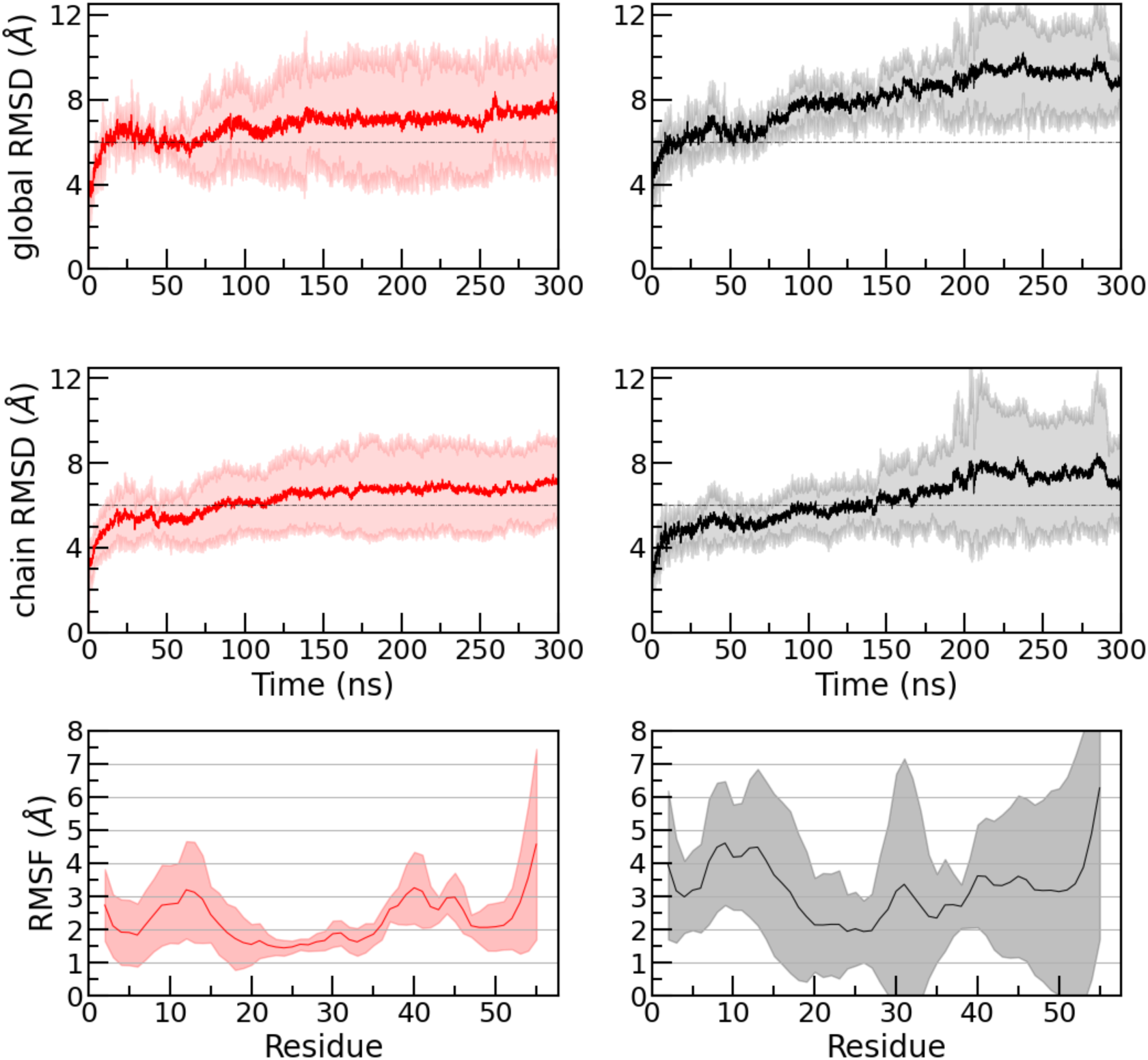
Evolution of the global and chain RMSD measured in simulations of the fibril in absence (in black) and presence of SK9 (in red). Chain RMSD values are averages over all four chains in the SAA fibril and all three trajectories. Backbone RMSF values are for the final 200 ns of each trajectory. The shaded region represents the standard deviation. The RMSD and RMSF values are calculated with respect to the experimentally solved structure considering all backbone atoms in residues 2-55

The observed stabilization of the chain architecture is interesting as the amyloidogenic segment is binding to the outside of the chains, i.e., the stabilization has to be indirect by either enhancing stacking or packing of chains. Analyzing the contact pattern, we find that presence of SK9 leads to an increase in hydrogen bonds and hydrophobic contacts involved in the stacking of chains.

While at the same time the SK9 segment reduces the contacts and hydrogen bonds involved in the packing of the two folds, this loss of contacts is compensated by the new interactions with the SK9- segment. The binding probabilities of SK9- segments towards the SAA fibril shown in **Figure 13** indicate that the SK9- segment binds to the hydrophobic and aromatic residues in the N-terminal and the C-terminal region. The binding pattern, indicated by Figure 13, stabilizes residue-residue stacking contacts, but also reduces the inter-strand packing distance (shown in **Table 2**) through forming contacts with certain charged (E^26^ and D^33^), polar residues (N^27^) and hydrophobic residues (M^24^ and A^30^) at the packing interface. Here, we define the inter-strand packing distance by the center of mass distance between two folds at their packing interface (residues 28-31). Note that the data for the control differ slightly from the one listed in Ref. 17 as our simulations are with 300 ns longer than the 100 ns of the simulations of Ref. 17.

**Figure 13:**
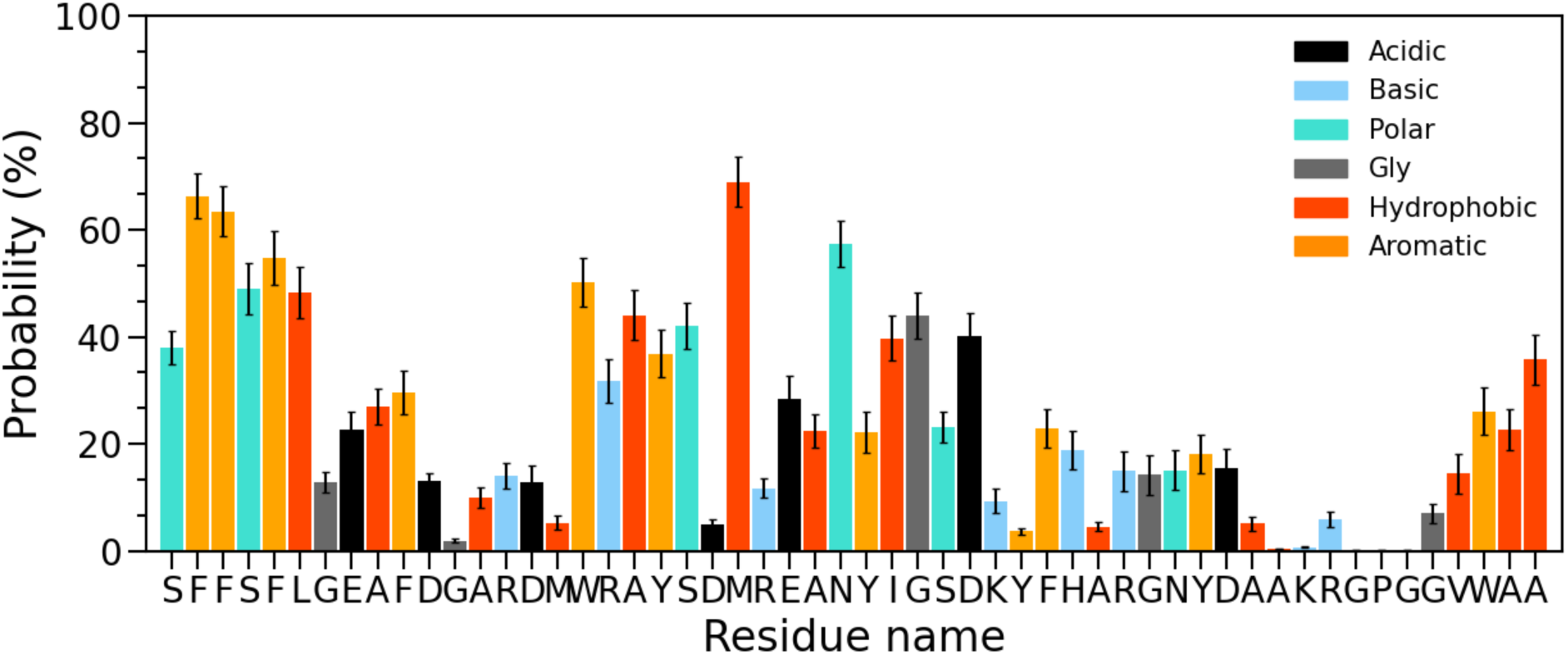
Residue-wise normalized binding probability of SK9- segment for the SAA fibril. Data were averaged over the final 200 ns of each trajectory.

**Table 2:** Mean values of stacking contacts and stacking hydrogen bonds; and of packing contacts and hydrogen bonds, averaged over all four chains in the SAA fibril and the final 200 ns of each trajectory. The corresponding standard deviations of the means are listed in brackets.

| System | stacking contact | stacking hydrogen-bonds | packing contact | packing hydrogen-bonds | Inter-stand packing distance (Å) |
| --- | --- | --- | --- | --- | --- |
| w/o SK9 | 624 (67) | 18 (4) | 83 (19) | 3.7 (1.7) | 9.9 (0.9) |
| with SK9 | 802 (66) | 25 (4) | 62 (24) | 2.0 (1.2) | 9.5 (0.5) |

In **Figure 14**, we plot the residue-residue stacking contact probability maps of the SAA fibril in the presence **(a)** or absence **(b)** of the SK9- segment. The difference between the two cases is shown in **Figure 14c**. Our data indicate that the residue-residue contact probability of the first 21 residues in the N-terminal region, the most hydrophobic and amyloidogenic segment of the protein, increases in presence of SK9. Especially enhanced are the π -π stacking and the hydrophobic interaction involving residues F^3^, F^4^, L^7^, F^6^, F^11^, A^10^, W^18^ and M^17^, and the propensity of residues R^15^ and D^16^ to form salt bridges. Additional stabilization of the fibril occurs at the C-terminal cavity (residues 23–51) through salt-bridges (D^23^-R^25^, R^25^-E^26^ and D^33^-K^34^), π -π stacking interactions (W^53^-W^53^) and hydrophobic interactions (A^54^-A^55^, A^55^-A^55^, A^54^-A^54^, W^53^-A^55^ and W^53^-A^27^).

**Figure 14:**
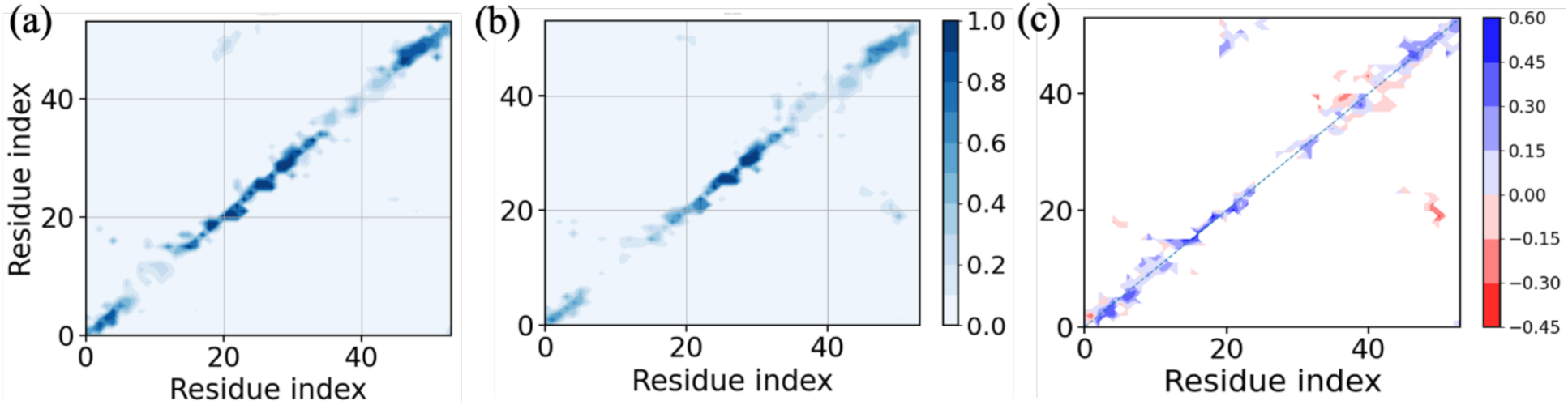
Residue-wise inter-layer contact probabilities measure in simulations of the two-fold two-layer SAA fibrils in the presence (a) and absence of (b) SK9- segment. A pair of residues is in contact if the center of mass distance is less than 7 Å. Data are averaged over the final 200 ns of each trajectory. The difference between the two stacking contact probabilities is shown in (c).

## Conclusions

The concentration of Human Serum Amyloid A (SAA) in acute COVID-19 patients can grow to levels that in patients with certain cancers or inflammatory diseases may cause systemic amyloidosis as a secondary illness. Hence, SARS-COV-2 infections may also increase the risk for SAA amyloid formation and subsequent pathologies. However, overexpression of SAA does not always lead to systemic amyloidosis, and mechanisms exist for downregulating SAA concentration and minimizing the risk for amyloidosis. In the present paper we use molecular dynamics to study how presence of SARS-COV-2 proteins may interfere with these protection mechanisms by changing the propensity for forming SAA amyloids. In order to reduce computational cost, we have restricted ourselves the nine-residue-segment S^55^FYVYSRVK^63^ (SK9) on the C-terminal tail of the SARS-COV2-Envelope protein whose location makes it likely to interact with SAA proteins.

Our simulations show that SARS-COV-2 proteins can increase the risk for SAA fibril formation by three mechanisms. First, binding of the SK9 reduces the stability of the biologically active SAA hexamer in which SAA transports lipids during inflammation, shifting the equilibrium toward monomers. As monomers are SAA proteins subject to enzymatic cleavage into smaller fragments, and only these fragments are found in SAA fibrils. Hence, by shifting the equilibrium toward the monomers, presence of the viral protein segment SK9 increases the risk for fibril formation. This risk is further enhanced by the interaction of SK9 with the SAA fragments, which increase the frequency of the aggregation prone form (called by us helix-weakened) and for this motif raises the propensity to form *β*-strands, especially for the first eleven residues known to be crucial for fibril formation. Finally, presence of the amyloidogenic segment SK9 also stabilizes SAA fibrils, moving further the equilibrium toward the fibril, and therefore enhancing the probability for amyloid formation. Hence, our simulations strengthen our hypothesis that SARSCOV-2 infections raise the risk for SAA amyloidosis during or after COVID-19. As SAA amyloidosis is characterized by formation and deposition of SAA amyloids in the blood vessels, causing inflammation and thrombosis, it may be behind the broad spectrum of severe and at times life-threatening cardiovascular, gastrointestinal, dermatologic, and neurological symptoms, commonly summarized as multisystem inflammatory syndrome (MISC) sometimes observed in COVID-19 survivors^11^.

It is interesting to speculate why amyloidogenic regions such as SK9 on SARS-COV-2 proteins seem to have such a pronounced effect on SAA amyloid formation. One possibility would be that fibril formation is part of the immune response and serves as a way to entrap and neutralize the virus. Such a microbial protection hypothesis^39, 40^ has been suggest in context of Herpes Simplex I infections and the development of Alzheimer’s Disease. Amyloid formation by SAA may serve a similar role, with SAA amyloidosis would be a consequence of this mechanism becoming overwhelmed. Further work will be needed to test this hypothesis.

## Author contributions

Conceptualization: AKJ, UH Methodology: AKJ, UH Investigation: AKJ, ABG, UH Visualization: AKJ, ABG Supervision: UH Writing—original draft: AKJ, UH Writing—review & editing: AKJ, ABG, UH

## Notes

The authors declare no competing financial interest.

## ACKNOWLEDGMENTS

The simulations in this work were done using the SCHOONER cluster of the University of Oklahoma, FRONTERA on TACC (under grant MCB20016), or XSEDE resources allocated under grant MCB160005 (National Science Foundation). We acknowledge financial support from the National Institutes of Health under research grant GM120634. We thanks Alan J. Ray for help with the figures.

